# Catabolic pathway acquisition by soil pseudomonads readily enables growth with salicyl alcohol but does not affect colonization of *Populus* roots

**DOI:** 10.1101/2024.01.17.575957

**Authors:** Stephan Christel, Alyssa A. Carrell, Leah H. Burdick, Manuel I. Villalobos Solis, Paul E. Abraham, Sara S. Jawdy, Julie E. Chaves, Nancy L. Engle, Timkhite-Kulu Berhane, Tao Yao, Jin-Gui Chen, Wellington Muchero, Timothy J. Tschaplinski, Melissa A. Cregger, Joshua K. Michener

## Abstract

Horizontal gene transfer (HGT) is a fundamental evolutionary process that plays a key role in bacterial evolution. The likelihood of a successful transfer event is expected to depend on the precise balance of costs and benefits resulting from pathway acquisition. Most experimental analyses of HGT have focused on phenotypes that have large fitness benefits under appropriate selective conditions, such as antibiotic resistance. However, many examples of HGT involve phenotypes that are predicted to provide smaller benefits, such as the ability to catabolize additional carbon sources. We have experimentally reproduced one such HGT event in the laboratory, studying the effects of transferring a pathway for catabolism of the plant-derived aromatic compound salicyl alcohol into soil isolates from the *Pseudomonas* genus. We find that pathway acquisition enables rapid catabolism of salicyl alcohol with only minor disruptions to existing metabolic and regulatory networks of the new host. However, this new catabolic potential does not confer a measurable fitness advantage during competitive growth in the rhizosphere. We conclude that the phenotype of salicyl alcohol catabolism is readily transferred by HGT but is selectively neutral under environmentally-relevant conditions. We propose that this condition is common and that HGT of many pathways will be self-limiting, because the selective benefits are small and negative frequency-dependent.

## INTRODUCTION

Due to competing processes of gene gain by horizontal gene transfer (HGT) and gene loss, bacterial gene content can vary widely even between strains of the same species [1, 2]. These ‘accessory’ genes, which are only present in a subset of strains, often alter the potential niche of the host [3], for example by encoding pathways to assimilate additional nutrients or tolerate new stresses [4, 5]. However, the fitness effects of accessory pathway acquisition are unclear, with arguments both for models that are largely adaptive or largely neutral [6, 7].

Any such discussion of fitness effects must include both the benefits of pathway acquisition as well as the associated costs [8]. The fitness effect of an accessory gene depends on a broad range of epistatic interactions, both with its host and the environment [9]. For example, acquisition of a pathway that provides access to a new niche will only be beneficial if the costs of pathway acquisition and integration are low while the benefits of niche expansion are high. These costs and benefits will depend on details of the genome content of the host strain and the biotic and abiotic conditions of the environment that the host inhabits.

The costs and benefits from HGT are often analyzed through knockout studies, removing putative horizontally-transferred genes and measuring changes in phenotype or growth [10]. However, these experiments are confounded by historical evolution following gene transfer, which can mitigate the costs of newly-acquired genes or introduce new dependencies [11, 12]. The effects of HGT can be more directly assessed by targeted pathway transfer in the laboratory followed by analysis of changes in phenotype and fitness [13].

If the benefits of HGT outweigh the costs, then targeted pathway transfer provides a potential opportunity to deliberately manipulate bacterial colonization, with applications in health, agriculture, and environmental remediation. For example, generating a new niche by feeding a marine polysaccharide to mice allowed specific colonization by a bacterium that had been engineered to contain the associated catabolic pathway [14]. In this case, the costs of pathway acquisition were low and the benefits were high. In general, this situation is likely to be rare, as many such ecological niches will already be filled by native microbes [15].

Even when an open niche is available, the utility of gaining access to a new niche may be small if the host already has access to other niches. For example, a recalcitrant environmental pollutant is an open niche that could be exploited as a carbon and/or energy source. However, when introducing allopatric microbes for bioremediation, metabolic specialists are more successful than generalists, likely because they have fewer alternative niches available in their new environment [16, 17].

Similar to the gut and soil, the rhizosphere is a complex environment with abundant metabolic niches and opportunities for HGT [18]. Plant root exudates provide diverse carbon sources that support high microbial populations [19]. Metabolic cross feeding and spatial structure further broaden the range of available niches [20]. These factors are particularly significant in perennial plants that can maintain a dynamic microbiome across multiple seasons [21].

Poplar (*Populus* sp.) trees provide a tractable model system to study microbial dynamics in the perennial rhizosphere [22]. Poplar trees exude large quantities of phenolic compounds derived from salicyl alcohol (SA), including salicin, populin, and tremuloidin [23]. These compounds are thought to act primarily as inhibitors of herbivory [24] but also serve as potential carbon sources for soil microbes [25]. An intermediate in the SA catabolic pathway, salicylic acid is a major component of plant exudates and thought to influence microbiome community composition [26]. Therefore, SA is a representative microbial metabolic niche in the rhizosphere and HGT of pathways for SA catabolism is likely to occur frequently. The persistence of these pathways after transfer will depend on the costs and benefits of pathway acquisition. Pseudomonads are abundant members of the rhizosphere microbiome and known for their aromatic catabolic potential, making them likely donors and recipients of SA catabolic pathways [27, 28].

In this work, we have assessed the utility of acquiring a pathway for catabolism of SA (Figure 1A). We show that salicyl alcohol catabolism is common among strains isolated from the *Populus* rhizosphere and this phenotype can readily be transferred into strains that do not natively possess it. The fitness costs and physiological disruption due to pathway acquisition are small. However, root colonization assays show that the fitness benefits are also minimal, even under conditions designed to maximize these effects. We conclude that acquisition of this catabolic pathway is functionally beneficial, in that it provides new capabilities at minimal cost, but selectively neutral.

**Figure 1:**
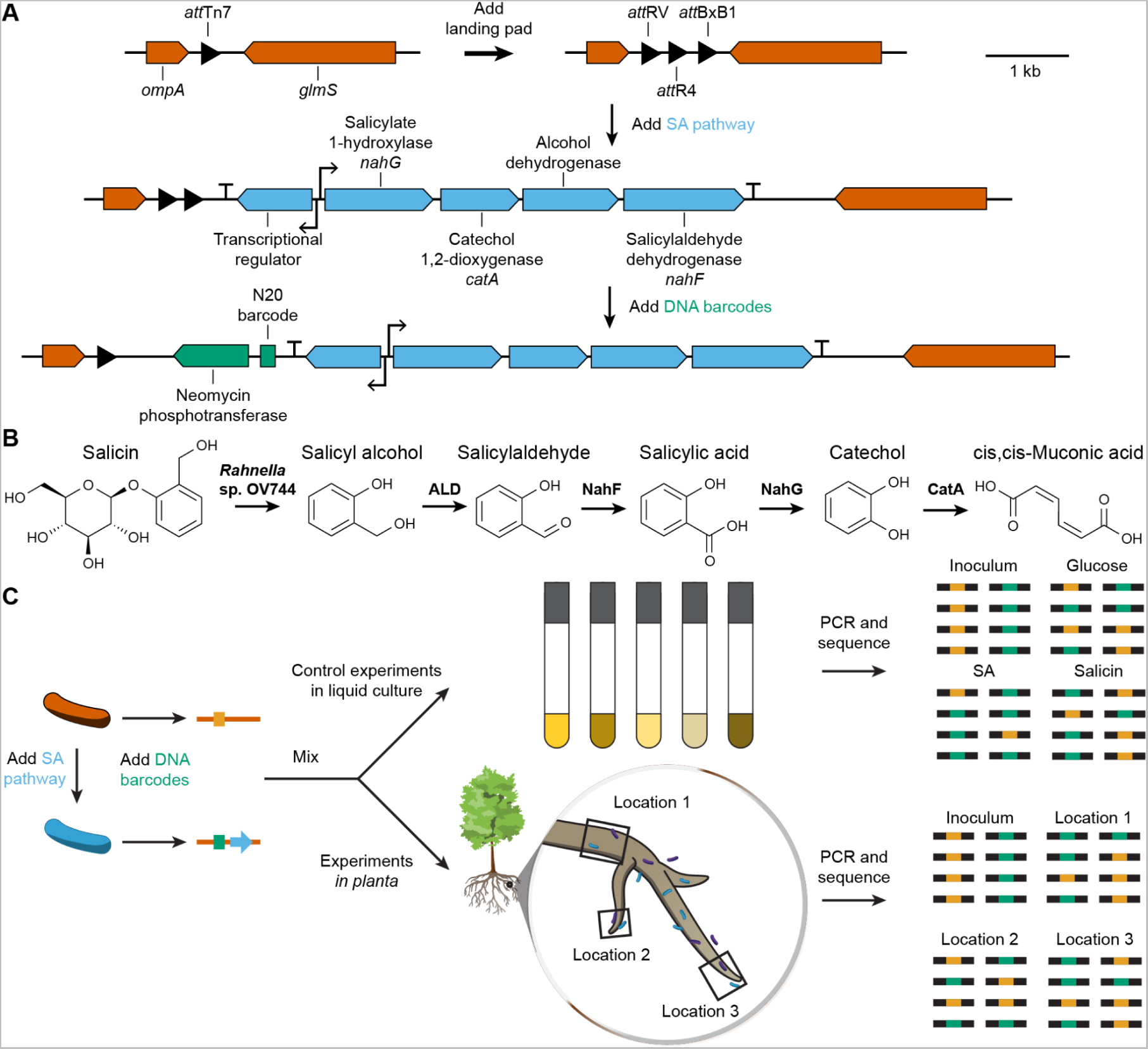
(A) To mimic HGT, the salicyl alcohol (SA) catabolic pathway from *Pseudomonas* sp. GM16 was transferred to the Tn7 *att* site in other *Pseudomonas* strains. DNA barcodes were then introduced to allow strain tracking *in situ*. (B) The proposed pathway for salicin degradation is initiated by a glycosyltransferase in a complementary strain, such as *Rahnella* sp. OV744, followed by successive oxidation to *cis*,*cis*-muconic acid. The tested *Pseudomonas* strains contain native pathways for assimilation of muconic acid. (C) Barcode amplicon sequencing was used to measure fitness effects of pathway acquisition, both in liquid culture and in the rhizosphere.

## RESULTS AND DISCUSSION

### Pseudomonad growth with salicyl alcohol

To evaluate the frequency of SA catabolism in rhizosphere pseudomonads, we tested for growth with SA among nine diverse *Pseudomonas* strains previously isolated from *Populus* roots (Figure 2A) [29]. Eight of the isolates grew in M9 minimal medium with SA as the sole source of carbon and energy, including the previously-characterized *Pseudomonas* sp. GM16 (hereafter ‘GM16’) [25]. One strain, *Pseudomonas* sp. GM17 (hereafter ‘GM17’), did not grow under these conditions (Figure 2B).

**Figure 2:**
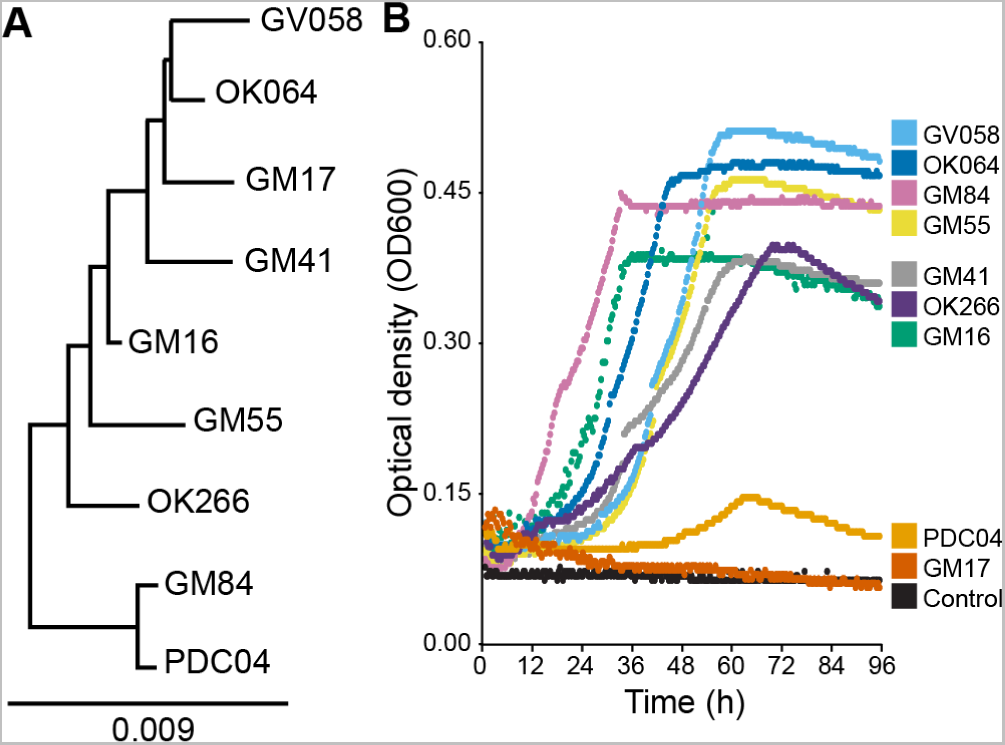
(A) 16S phylogenetic tree of *Pseudomonas* strains isolated from *Populus* and tested for SA catabolism. *Cellovibrio japonicus* was used as an outgroup (not shown). (B) Growth curves of *Pseudomonas* strains in M9 minimal medium with salicyl alcohol as the sole source of carbon and energy. One representative curve is shown for each strain, chosen from three biological replicates.

Given the high frequency of SA catabolism in these isolates, we sought to understand the factors limiting the dissemination or retention of this pathway in GM17. We initially hypothesized that deleterious interactions between a newly-introduced SA pathway and the native metabolic pathways of the potential hosts would prevent successful transfer into GM17 [30, 31]. To test this hypothesis, we engineered the SA catabolic pathway into several *Pseudomonas* isolates and measured changes in catabolic activity. We chose GM17 as a representative non-catabolizing soil isolate, *Pseudomonas putida* KT2440 as a non-catabolizing laboratory reference strain, and *Pseudomonas* sp. PDC04 as a representative poorly-catabolizing isolate.

We first integrated *attP* sites for heterologous serine integrases into the T7 phage integrase *att* site in each recipient strain [32–34]. We then used the heterologous BxB1 *attP* site to stably introduce the SA catabolic pathway from GM16, including its putative SalR regulator, into the genomes of the recipients. When we measured growth with SA, we found that all three engineered strains could now grow with SA (Figure 3A). Strain PDC04, which could naturally metabolize SA, grew more rapidly with SA after introduction of the heterologous catabolic pathway. We did not observe noticeable changes in growth with glucose between the wildtype and engineered strains (Figure 3B).

**Figure 3:**
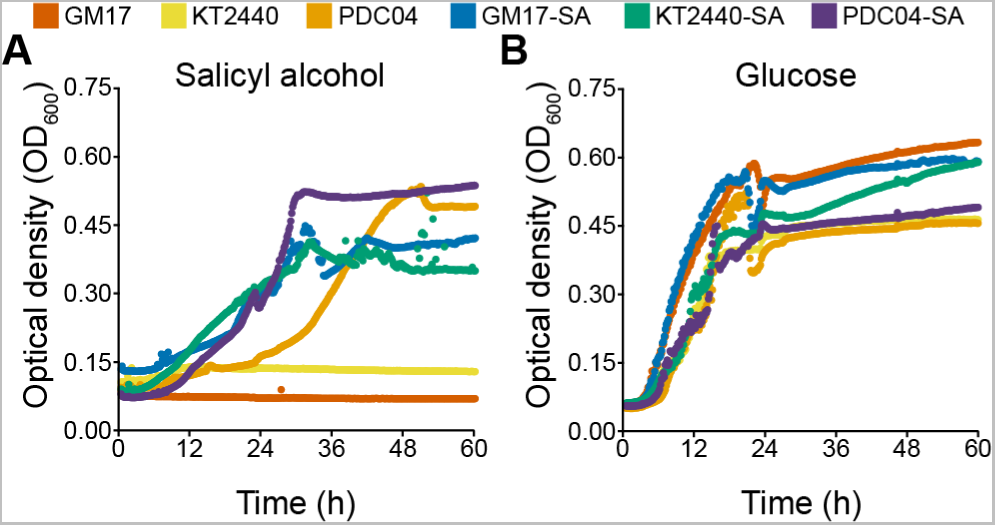
Growth of wildtype and engineered *Pseudomonas* strains. The ‘SA’ prefix indicates that the strain contains a genomically-integrated SA catabolic pathway. Individual strains were grown in M9 minimal medium containing 1 g/L glucose or salicyl alcohol as the sole carbon and energy source.

### Proteomics analysis of pathway integration

While the engineered strains grew readily with SA, we hypothesized that pathway acquisition might impose subtle stresses on the new host that would limit pathway retention under more stringent selective conditions [35, 36]. To test this hypothesis, we performed global proteomic analysis of one wild-type strain, GM16, and three engineered strains, GM17-SA, PDC04-SA, and KT2440-SA. Each strain was grown with SA as the sole carbon source and compared to the corresponding strain grown with glucose.

On average, 27.7±2.6% of measured proteins were detected at significantly (q<0.05 and log_2_ fold change>2) higher or lower abundance during growth with SA. This broad shift in expression could indicate a relatively large physiological perturbation. However, based on gene annotations, the vast majority of differentially expressed proteins were likely directly involved in the degradation of SA and its cascading products.

We used OrthoMCL to group and compare protein function rather than often-misleading sequence identity. This analysis revealed that all but a few significantly differentially abundant proteins could be placed along the flow of carbon from SA to acetyl/succinyl-CoA metabolism and TCA cycle (Figure 4) and that this behavior was indeed shared between all three SA strains plus the original host of the SA degradation pathway, GM16. Very few ortholog groups shared differential expression patterns in all engineered strains but not GM16. Among them was a hydroxybenzoate monooxygenase (Orthogroup OG6_152207), a glutaryl-CoA dehydrogenase (OG6_101810), as well as several dehydrogenases, carboxylases, and ligases involved in CoA and pyruvate metabolism (Figure 4). The lack of such a distinct ‘mutant signature’, i.e., the absence of a large set of differentially abundant proteins common exclusively to engineered strains, indicated that the introduced pathway did not disrupt genetic regulation in its new host strains. Furthermore, detailed abundance analysis of the four proteins newly introduced into the engineered strains revealed that they were only minimally expressed during growth on glucose (data not shown), confirming that the regulatory systems from GM16 also remained functional. Based on the proteomics results, we concluded that the pathway is active, properly regulated, and does not cause significant stresses to the new host bacteria.

**Figure 4:**
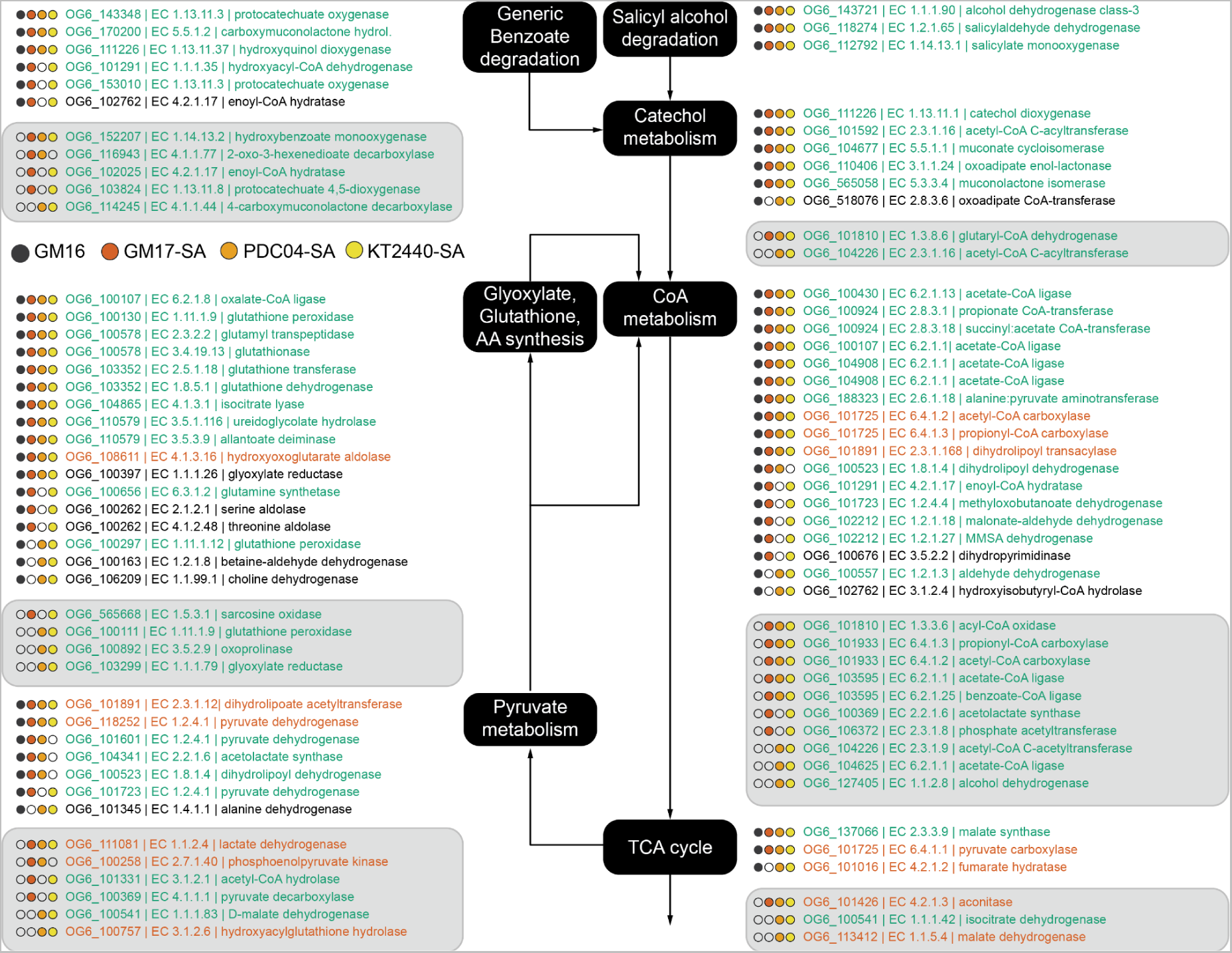
Model of proteins with significantly increased abundance in wildtype GM16 and engineered GM17, PDC04, and KT2440 strains when grown with SA compared to glucose. Filled circles indicate that the orthologous group was identified to be significantly differentially expressed in the corresponding strain during growth with SA versus glucose. Orthologous groups were grouped according to their predicted function and displayed along the pathway of SA oxidation (black boxes). Green text indicates increased expression in presence of SA, red text decreased expression, and black text contradicting expression in the strains. Light gray background indicates orthologous groups uniquely identified in the mutants.

### Impact of pathway acquisition on root colonization

Since the SA pathway was functional and minimally disruptive, we next tested its effect under more environmentally-realistic conditions, during colonization of *Populus trichocarpa* roots. To track changes in relative bacterial abundance, we introduced random 20 nt DNA barcodes flanked by conserved primer binding sites into the R4 *attB* sites in wild-type and engineered GM17 (Figure 1A). We downsampled each library to approximately 10,000 barcodes per strain and sequenced each library to identify the barcodes that uniquely identified each strain. Changes in relative abundance of the two strains can then be tracked by targeted amplicon sequencing of the barcode region.

To test the accuracy of the assay, we mixed the barcoded populations of wild-type and engineered GM17 and grew the mixed culture in liquid culture with glucose or salicyl alcohol as the sole carbon source. We sequenced amplicons from the inoculum and saturated cultures and determined changes in relative abundance of the wild-type and mutant strains (Figure S1). We observed no change in relative abundance of the SA mutant after growth with glucose, but an enrichment for the SA mutant after growth with SA. These results are consistent with prior growth experiments using pure cultures, showing that the SA pathway is active and provides a growth advantage when SA is the sole carbon and energy source. To determine the sensitivity of this assay, we also used the mixed culture to inoculate roots of tissue-cultured *Populus trichocarpa* grown in sterile clay. We grew the resulting plants for 21-28 days before harvesting the trees. We then dissected the roots into segments ranging in mass from approximately 10 mg to less than 0.1 mg and performed amplicon sequencing on the barcodes (Figures S2+S3). We reliably amplified barcodes from root segments with masses less than 1 mg.

Next, we inoculated tissue-cultured *P. tremula* x *P. alba* ‘INRA 717-1B4’ with the GM17 mixed culture and measured changes in relative abundance. We hypothesized that, if the SA pathway provided a fitness advantage to its host during colonization, then the abundance of barcodes from the engineered strain would increase relative to the wild-type strain (Figure 1C). Since plant metabolite profiles are expected to vary spatially based on root architecture, we tested replicate samples from primary, secondary, and tertiary root segments, root hairs, and root tips (Figure 5A). However, we observed no significant differences in abundance in any location (Figure 5B). We concluded that the presence of the SA pathway was providing a minimal net fitness benefit, either because the gross benefits were small or because there was a corresponding cost to SA pathway maintenance and expression.

**Figure 5:**
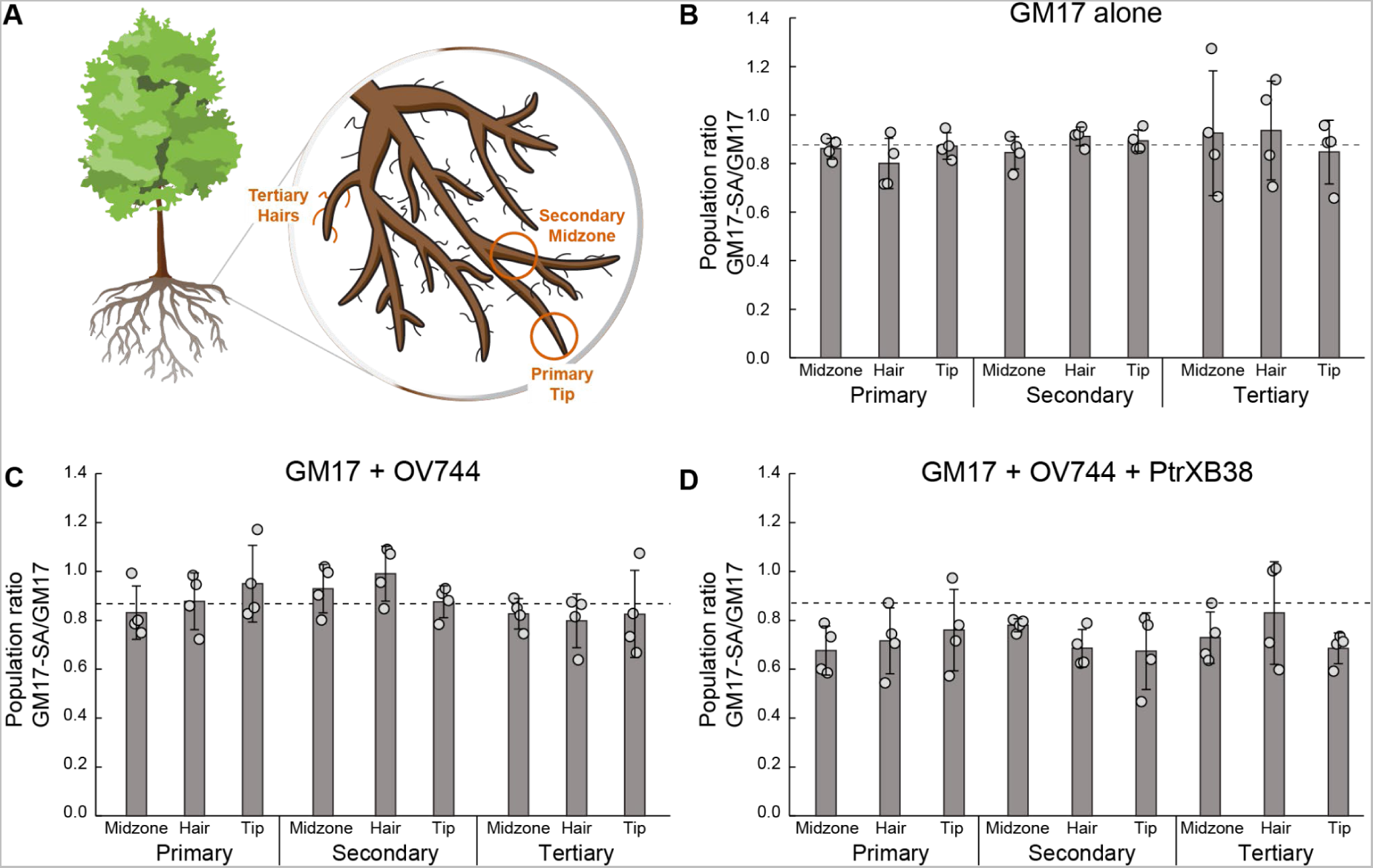
SA catabolism provides no fitness effect during plant colonization. (A) Mixtures of bacteria were inoculated onto axenic *P. tremula* x *P. alba* ‘INRA 717-1B4’ cuttings and grown for 28 days. Population ratios were sampled at a range of sites, including primary/secondary/tertiary roots at the tip/midzone/root hairs. Representative sites are shown in red. (B) A mixture of wild-type and engineered GM17 was inoculated onto axenic *P. tremula* x *P. alba* ‘INRA 717-1B4’. The population ratios before and after cultivation were determined by barcode amplicon sequencing. The dashed line shows the population ratio of the inoculum. Error bars show one standard deviation, calculated from the four biological replicates shown. (C) Same as B, but with the addition of the salicin-degrading bacterium *Rahnella* sp. OV744. (D) Same as C, but using an salicin-overproducing *PtrXB38-OE* line of *P. tremula* x *P. alba* ‘INRA 717-1B4’.

### Effects of epistatic interactions that benefit SA catabolizers

While *Populus* secretes small amounts of SA, it primarily secretes SA conjugates, including salicin (Figure S4). Neither wild-type nor engineered GM17 can degrade salicin, which might limit the effect of SA catabolism in the absence of an accessory deglycosylation pathway. Since co-cultures of GM16 and *Rahnella* sp. OV744 (hereafter ‘OV744’) have been shown to fully degrade salicin, we tested whether the presence of OV744 would alter the fitness effect of SA catabolism in GM17. We first repeated the *in vitro* assays, inoculating wild-type and engineered GM17 into M9 minimal medium with salicin as the sole carbon source in the presence and absence of OV744. Using amplicon sequencing to specifically track changes in abundance of wild-type and engineered GM17 from the mixed culture, we observed that the SA pathway provided a fitness benefit during growth with salicin only when OV744 was also present in the culture (Figure S5). These results are consistent with the model that OV744 converts salicin into SA, which is then available for catabolism by the GM17 SA mutant (Figure 1B).

We repeated the *P. tremula* x *P. alba* ‘INRA 717-1B4’ inoculations using the GM17 mixed library in the presence of OV744 and sampled a range of root positions after 28 days of growth. We again observed no change in the relative abundance of the wild-type versus engineered GM17 strains (Figure 5C). By comparison to the control experiments, we conclude that SA production from salicin by OV744 was too low to provide a fitness benefit to the GM17 SA mutant.

To further amplify the potential benefits of SA catabolism, we repeated these experiments using a *P. tremula x P. alba* INRA 717-1B4 mutant that overexpresses the *PtrXB38* gene [37]. This mutant has been shown to produce significantly higher concentrations of salicin in the roots (Figure S4). However, as in our previous experiments, we saw no enrichment for the SA-degrading GM17 strain during growth with the *PtrXB38-OE* plants either in the presence or absence of OV744 (Figure 5D and Figure S5). The salicin concentration is higher in the *PtrXB38-OE* plants, which would presumably provide more SA after hydrolysis by OV744. However, the concentration may still be too low compared to the other available carbon sources to provide a significant fitness difference.

## CONCLUSIONS

In combination, our results demonstrate that acquisition of a pathway for SA catabolism, either in the laboratory through metabolic engineering or presumably in nature through HGT, can readily provide rhizosphere isolates with new metabolic capabilities. The pathway for SA catabolism imposes minimal disruption on the native pathways of the new host bacterium. However, SA catabolism did not provide a fitness benefit during root colonization, even under conditions that are designed to favor SA catabolizers.

We hypothesize that, under the conditions tested, GM17 can access sufficient metabolic niches that the availability of a new niche does not alter its fitness. In a more realistic microbial community, it might face additional competition in these other niches and gain a larger proportional advantage from SA catabolism. However, as we have shown, a more realistic community would also contain many other SA-catabolizing microbes, so the benefits of the new niche would also be smaller.

In general, our results are consistent with a model where catabolic pathways spread through HGT until the benefits of pathway acquisition decline to the point that they are balanced by the small costs of pathway maintenance. While catabolic potential can drive significant changes in colonization, we suggest that those examples will be limited to rare metabolites that are not efficiently exploited by the native microbiota and specialist microbes that have few alternative niches available. These conclusions are consistent with prior evidence from bioaugmentation for bioremediation and offer cautionary guidance for efforts to control colonization of introduced microbes into native communities through engineering carbon utilization.

## MATERIALS AND METHODS

### Strains

*Pseudomonas* strains used in this study were *Populus*-derived isolates from the ORNL Plant-Microbe Interfaces strain collection [29]. *P. putida* KT2440 was acquired from the American Type Culture Collection. Genome sequences (complete or partial) are available for all used wild type strains and can be retrieved from online repositories (Table 2).

**Table 1:** Differentially abundant proteins identified in analysis of four wild-type/SA strain pairs grown in triplicate in presence of 0.1% glucose versus 0.1% salicyl alcohol.

| Strain | GM16 | KT2440 | GM17 | PDC04 |
| --- | --- | --- | --- | --- |
| **Total detected proteins** | 2069 | 1891 | 2070 | 1903 |
| **Upregulated Glucose** | 217 | 238 | 172 | 245 |
| **Upregulated SA** | 277 | 351 | 389 | 303 |
| **Total differentially abundant proteins** | 494 | 589 | 561 | 548 |
| **Percentage of total** | 23.9 | 31.1 | 27.1 | 28.8 |

\*p<0.05 and log2fold-change>2

**Table 2:** Strains used in this study.

| Strain | Genotype | Reference |
| --- | --- | --- |
| *Pseudomonas* sp. GM16 | Wildtype | Carper *et al.* |
| *Pseudomonas* sp. GM17 | Wildtype | Carper *et al.* |
| *Pseudomonas* sp. PDC04 | Wildtype | Carper *et al.* |
| *Pseudomonas putida* KT2440 | Wildtype | ATCC |
| *Rahnella* sp. OV744 | Wildtype | Carper *et al.* |
| *Pseudomonas* sp. GM41 | Wildtype | Carper *et al.* |
| *Pseudomonas* sp. GM55 | Wildtype | Carper *et al.* |
| *Pseudomonas* sp. GM84 | Wildtype | Carper *et al.* |
| *Pseudomonas* sp. GV054 | Wildtype | Carper *et al.* |
| *Pseudomonas* sp. GV058 | Wildtype | Carper *et al.* |
| *Pseudomonas* sp. OK064 | Wildtype | Carper *et al.* |
| *Pseudomonas* sp. OK266 | Wildtype | Carper *et al.* |
| JMP43 | GM16 *attTn7::attBxB1/attRV/attR4* | This work |
| JMP44 | GM17 *attTn7::attBxB1/attRV/attR4* | This work |
| JMP52 | PDC04 *attTn7::attBxB1/attRV/attR4* | This work |
| JMN42 | KT2440 *attTn7::attBxB1/attRV/attR4* | This work |
| JMP88 | GM17 *attTn7::(attBxB1::SA-Reg)/attRV/attR4* | This work |
| JMP90 | PDC04 *attTn7::(attBxB1::SA-Reg)/attRV/attR4* | This work |
| JMP91 | KT2440 *attTn7::(attBxB1::SA-Reg)/attRV/attR4* | This work |
| JMP96 | GM17 *attTn7::attBxB1/attRV/(attR4::Barcode s)* | This work |
| JMP97 | GM17 *attTn7::(attBxB1::SA-Reg)/attRV/(attR4::Barcodes)* | This work |

### Construction of a broad host range tri-*attP* landing pad

A landing pad containing three phage integrase *attP* recognition sequences was designed for chromosomal integration using a broad host range mini-Tn7 vector method [32]. First, the tri-*attP* landing pad sequence was cloned into a mini-Tn7 vector, followed by co-transformation into each *Pseudomonas* strain with a plasmid encoding Tn7 transposition pathway expression. Finally, the antibiotic selection marker for the mini-Tn7 vector was removed by Flp-mediated excision. The mini-Tn7 vector (pUC18T-mini-Tn7T-GM, Addgene plasmid # 63121), Tn7 transposase expression plasmid (pTNS, Addgene plasmid # 64967), and the FLP recombinase expression plasmid (pFLP3, Addgene plasmid # 64946) were gifts from Herbert Schweizer [38].

The R4, Bxb1 and RV phage integrase *attP* recognition sequences were designed into a single landing pad sequence and synthesized *de novo* (Twist Biosciences). The landing pad sequence was PCR amplified from the vector, and cloned into the mini-Tn7 vector following manufacturer instructions (NEBuilder HiFi Assembly Master Mix) to generate a mini-Tn7 vector + Landing Pad plasmid pJM442.

### Construction of *Pseudomonas* recipient strains

Each wildtype *Pseudomonas* strain (Table 2) was transformed with pJM442 plasmid using quad-parental conjugation as previously described [32]. *E coli* WM6062, a diaminopimelate (DAP) auxotroph strain carried the mini-Tn7 vector + Landing Pad plasmid (pJM442), *E. coli* Pir1 carried the pTNS2 Tn7 transposase expression plasmid, and *E. coli* DH5α carried a plasmid containing the conjugation machinery (pRK2073_Kan^R^). All four strains were grown overnight in LB medium supplemented with the appropriate antibiotics or nutrients (DAP), at 30°C (*Pseudomonas*) or 37°C (*E. coli*). Saturated cultures were combined into a single Eppendorf tube at equal 100 mL volumes. The mixed culture was then centrifuged at room temperature at 7000 x *g*, washed twice in 1 mL of sterile 10 mM MgSO_4_, and resuspended into 30 uL 10 mM MgSO_4_. The final resuspension was dropped onto a pre-dried LB agar plate supplemented with DAP (60 mg/mL), and incubated overnight at 30°C. The next day, the cells biomass was scraped from the agar plate, resuspended into 5 mL sterile 10 mM MgSO_4_, and serially diluted. Then, 100 µL of the 10^-3^ and 10^-5^ dilutions were spread onto LB agar plates supplemented with 100 µg/mL gentamicin without DAP and incubated at 30° C overnight or until clear colonies appeared. Proper integration of the landing pad into each strain was verified by whole genome resequencing.

To complete the recipient strain construction, the gentamycin resistance cassette was removed from each strain using Flp-mediated excision. Chemically competent versions of each strain were generated [39]. Single colonies of each strain were used to inoculate LB medium supplemented with 100 µg/mL gentamycin and grown to saturation overnight. Saturated cultures (1 mL) were transferred to pre-chilled Eppendorf tubes and centrifuged at room temperature, for 1 minute at 13,000g. The supernatant was decanted, and the pellet was resuspended and washed twice in 1mL cold 0.1 mM MgCl_2_ at room temperature. After the second wash, the pellet was resuspended with 1 mL cold TG salt (75 mM CaCl_2_, 6mM MgCl_2_, and 15% glycerol), incubated for 10 min on ice, centrifuged as above and resuspended with 200 µL TG salt. The cells were then flash frozen in liquid nitrogen, and kept at −80°C until used for transformation.

Aliquots of 100 µL of each chemically competent *Pseudomonas* strain were mixed with 100 ng of the FLP recombinase expression plasmid (pFLP3), incubated together on ice for 15 minutes, followed by a 2 min heat shock at 37 °C. Cells were immediately resuspended in 900 µL SOC and allowed to recover for 30 minutes at 30 °C, followed by plating onto LB agar supplemented with 25 µg/mL tetracycline, and incubation 30 °C overnight, or until clear colonies became visible. The absence of the gentamicin cassette was checked by patching single colonies simultaneously onto two LB agar plates, one containing 100 µg/mL gentamicin, or 25 µg/mL tetracycline. Both plates were incubated at 30 °C overnight or until colonies appeared. Successful transformants resulted in the formation of colonies that grew on LB with tetracycline, but did not grow in the presence of gentamycin.

The FLP recombinase expression plasmid pFLP3 was cured from the transformant cells using sucrose counter selection. Successful FLP recombinase transformants were streaked to single colony on YT-25 % sucrose agar (10 g/L yeast extract, 20 gl/L tryptone, 250 g/L sucrose, 18 g/L agar) for counter-selection against the FLP plasmid, incubating at 30 °C for ~30-48 hours. Surviving colonies were then re-streaked onto fresh YT-25% sucrose plates and incubated at 30°C for 16 hours to remove any enduring *sacB*-containing cells. Single colonies were simultaneously patched onto 2 different LB agar plates, one containing 25% sucrose, and one containing tetracycline. Successfully cured cells resulted in colonies that were able to grow in the presence of sucrose, but not tetracycline. The final colonies were further verified for successful removal of the gentamycin resistance cassette by whole genome resequencing.

### Salicyl alcohol degradation pathway design, synthesis, and integration

The SA degradation operon was amplified from *Pseudomonas* sp. GM16 genomic DNA using primers SA-Reg FWD and SA-Reg REV (Table 4) and polymerase Q5 (New England Biolabs, Massachusetts, USA). The destination plasmid pGW44 [33] was linearized using primers pGW44 FWD and pGW44 REV. The SA pathway was then introduced into pGW44 using the Gibson Assembly Cloning Kit (New England BioLabs), creating plasmid pJM455. The complete pathway was then genomically integrated into target strains using the BxB1 *attB* site, followed by removal of the kanamycin selection marker, as described previously [33].

**Table 3:** Plasmids used in this study.

| Plasmid | Genotype | Reference |
| --- | --- | --- |
| pUC18T-mini-Tn7T-GM |  | Choi et al., 2005<br>Addgene 63121 |
| pTNS2 |  | Choi et al., 2005<br>Addgene 64968 |
| pFLP3 |  | Choi et al., 2005<br>Addgene 64946 |
| pJM442 | pUC18T-mini-Tn7T-GM +<br>*attBR4/attBBxBI/attBRV* | This work |
| pGW44 | BxB1*attP colE1 kanR* | Elmore et al., 2023 |
| pJM455 | pGW44 + SA catabolism<br>pathway | This work |
| pJH207 | R4*attP colE1 kanR* | Elmore et al., 2023 |
| pJM488 | pJH207 + 20 nt barcodes | This work |

**Table 4:**
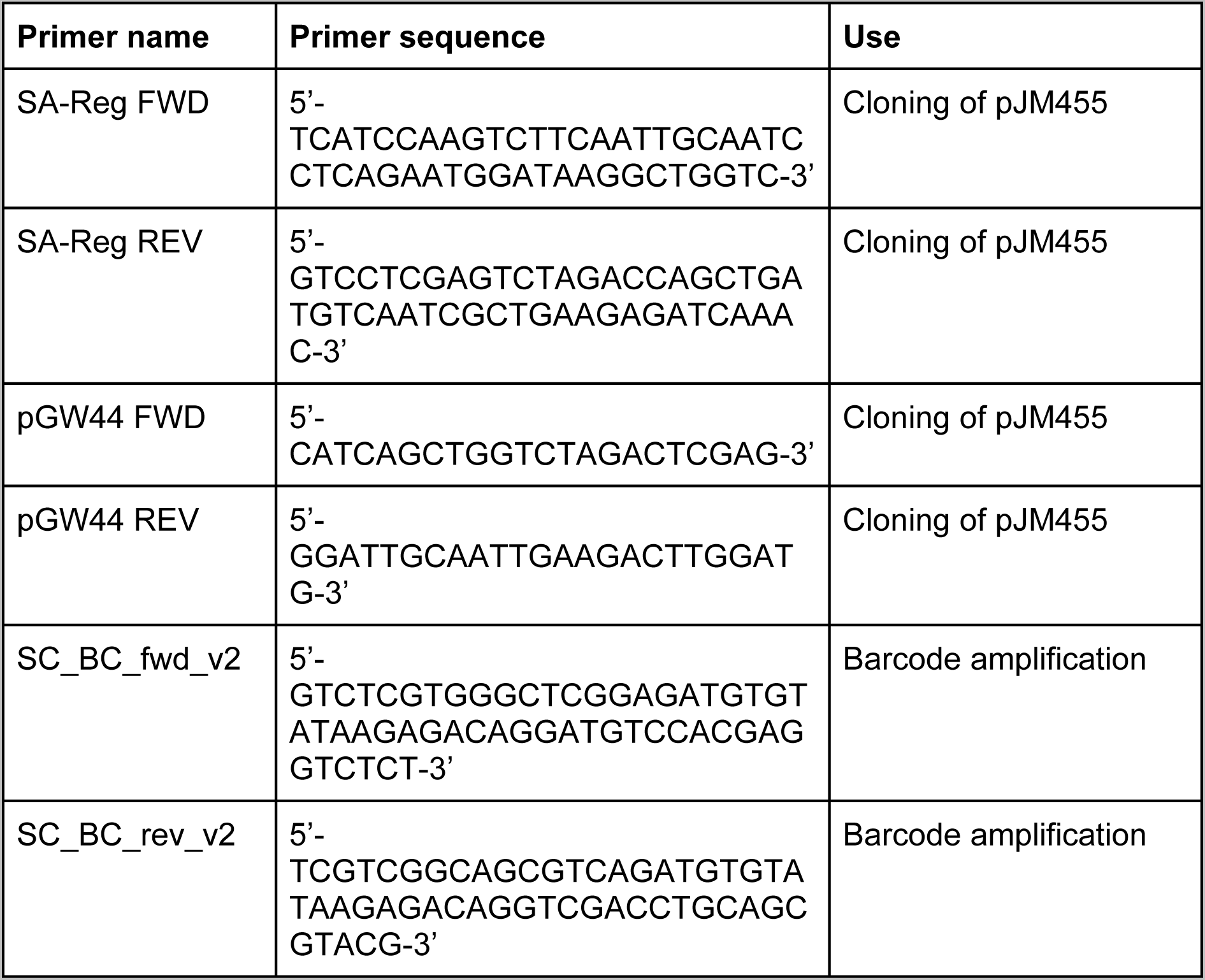
Primers used in this study.

### Growth analysis

To measure growth kinetics with specific carbon sources, strains were grown to saturation in M9 minimal medium with 1 g/L glucose as the sole carbon source. Saturated cultures were centrifuged for 3 minutes at 8000 x *g*, washed with M9 minimal medium without carbon, and then diluted 1:100 into 100 μL fresh medium containing the indicated carbon source. Cultures were grown at 30 °C in 96-well plates in an BioTek Epoch 2 shaking incubator (Agilent, Santa Clara, CA) for 72 hours.

### Wild type and mutant barcoding

To track and differentiate strains after eventual inoculation and growth on plant roots, a set of random DNA barcodes was also integrated into each genome. A plasmid library containing an R4 *attB* site and a stretch of 20 random nucleotides was synthesized (Biomatik, Ontario, Canada). The resulting barcode library was integrated into the R4 *attB* site of otherwise wild-type strains, containing only the landing pad, or strains with the SA catabolic pathway already integrated into the BxB1 *attP* site. The full libraries, which contained several million barcodes each, were subsampled to achieve an estimated library size of 10,000 barcodes per strain by spreading a dilution series on large LB agar plates and subsequently harvesting 10,000 colonies by washing a corresponding number of plates.

### Proteomics analysis

#### Cultivation

For proteomic analysis, wild type and mutant strains were grown in test tubes containing 10 ml M9 medium supplemented with 0.1 % glucose, 0.1% salicyl alcohol, or a combination of the two. A 5% salicyl alcohol stock solution was prepared in absolute ethanol and was added to the empty test tubes at the outset of media preparation, to allow evaporation of the ethanol before addition of the M9. Glucose overnight cultures of the strains were washed twice in M9 without an energy source, inoculated into the test tubes, and finally incubated under shaking (30 °C, 250 RPM) until reaching OD 0.5. Then, cells were harvested by centrifugation at 8000 × g for 5 min, supernatant was aspirated, and the cell pellet was immediately frozen at −80 °C until further processing.

#### Cell lysis and protein extraction and digestion

Cell pellets were solubilized with 500 µL of lysis buffer (4% sodium dodecyl sulfate (SDS) w/v in 100 mM Tris-HCl, pH 8.0). Samples were vortexed and then disrupted by bead beating for 5 mins with 0.15mm Zirconium oxide beads at 3:1 volume ratio of sample to beads. Samples were then placed in a heat-block for 10 min at 90°C. Approximately 400 µL of cell lysates were transferred to fresh Eppendorf tubes after centrifugation for 3 minutes at 21,000g.

Protein concentration was measured using a NanoDrop™ One^C^ instrument (Thermo Scientific). Each sample was adjusted to 10 mM dithiothreitol (DTT) and incubated at 90 °C for 10 minutes. Following DTT addition, samples were then adjusted to 30 mM iodoacetamide (IAA) to prevent reformation of disulfide bonds and incubated in the dark for 15 minutes. To isolate proteins, the protein aggregation capture method was employed [40]. Briefly, Ser-Mag beads and crude lysates were added to fresh Eppendorf tubes at a 1:1 protein to beads ratio, precipitated and captured by adjusting to 70% (v/v) LC-MS grade ACN, then washed with 1 mL ACN and 1mL of 70% LC-MS grade ethanol. Sequence-grade trypsin solution was added to a 1:75 (wt/wt) ratio of protein to trypsin and then additional Tris buffer was added to a final additional volume of 200 µL. Trypsin digestion was performed overnight at 37 °C under constant shaking at 600 rpm using an Eppendorf Thermomixer (Thermo Scientific). Proteins were digested a second time using the same protein to trypsin ratio of before, but this time incubated for 3 h at 37 °C under constant shaking at 600 rpm. After protein digestion, each sample was then adjusted to 0.5% formic acid (v/v) followed by vortexing and incubation at room temperature for 10 min. Each sample was then centrifuged at 21,000g for 10 min and supernatants transferred on top of pre-equilibrated 10 kDa MW cutoff Vivaspin 500 filters. Tryptic peptides flowthroughs were then collected after centrifugation at 12,000g for 10 min. Peptide concentrations were measured using the same Nanodrop instrument of before and transferred to autosampler vials for LC-MS/MS measurement.

#### LC-MS/MS

Peptide mixtures were analyzed using two-dimensional (2D) liquid chromatography (LC) on an Ultimate 3000 RSLCnano system (Thermo Fisher Scientific) coupled with a Q Exactive Plus mass spectrometer (Thermo Fisher Scientific). For each sample, aliquots equivalent to 2 µg of peptides were injected to an in-house built strong cation exchange (SCX) Luna trap column (5 µm, 150 µm X 50 mm; Phenomenex, USA) followed by a nanoEase symmetry reversed-phase (RP) C18 trap column (5 µm, 300 µm X 50 mm; Waters, USA) and washed with an aqueous solvent. Cellular peptide mixtures were separated and analyzed across one SCX fraction by eluting the peptides form the SCX column with a volume plug of 500 mM ammonium acetate followed by a 90-min organic gradient (250 nL/min flow rate) to separate peptides across an in-house pulled nanospray emitter analytical column (75 µm X 350 mm) packed with 35 cm of C18 Kinetex RP C18 resin (1.7 µm; Phenomenex, USA). Mass spectra were acquired with the Q Exactive Plus instrument in a top 10 data-dependent acquisition setup. MS spectra were collected within 300 to 1500 m/z with automatic gain control (AGC) target value of 3 × 10^6^ at a resolution of 70,000 with a maximum injection time (IT) of 25 ms. Precursor ions with charge states ≥2 and ≤ 5 and intensity threshold of 1.6 × 105 were isolated using a 1.6 m/z isolation width for higher-energy C-trap collision dissociation (HCD) with a normalized collision energy of 27 eV. MS/MS spectra were acquired at a resolution of 17,500 at m/z 200 with an AGC target value of 1 × 10^5^ and maximum IT of 50 ms. Dynamic exclusion was set to 20 s to avoid repeated sequencing of peptides. Each MS raw data file was processed by the SEQUEST HT database search algorithm and confidence in peptide-to-spectrum (PSM) matching was evaluated by Percolator [41] using the Proteome Discoverer v2.2 software. Peptides and PSMs were considered identified at q<0.01 and proteins were required to have at least one unique peptide sequence. Protein relative abundance values were calculated by summing together peptide extracted ion chromatograms. Protein abundances were normalized by LOESS and median central tendency procedures performed on log2-transformed values by InfernoRDN [42].

#### Ortholog analysis for cross-species comparison

Ortholog groups were constructed with OrthoMCL using pre-configured workflows at the VEuPathDB Galaxy site. In brief, all-versus-all BLASTP and the OrthoMCL algorithm were used to assign each organism-encoded protein to OrthoMCL groups (version OG6r1) with a 1e-05 expectation value cutoff for BLASTP and a 4 main inflation value for the clustering algorithm MCL.

### Metabolite analysis

Salicin and salicyl alcohol of the PtrXB38-OE and control plants were extracted from ~150 mg of frozen powdered root tissue twice overnight with 2.5 mL of 80% ethanol. Sorbitol (75 µL of 1mg/mL aqueous solution) was added to the first extract as an internal standard [37]. The two extracts were combined, and a 1 mL aliquot was dried under nitrogen for analysis. The dried extracts were silylated to produce trimethysilyl (TMS) derivatives by dissolving in 500 µL of silylation grade acetonitrile (Thermo Scientific, TS20062), followed by addition of 500 µL of N-methyl-N-trimethylsilyltrifluoroacetamide (MSTFA) with 1% trimethylchlorosilane (TMCS) (Thermo Scientific, TS48915) and heated for 1 h at 70 °C. After 2 days, 1 µL was injected into an Agilent Technologies 7890A GC coupled to a 5975C inert XL MS configured as previously described but with the following modification. Gas (He) flow was 1.20 mL per minute. Metabolite peaks were extracted using key mass-to-charge (m/z) selected ions to minimize interference with co-eluting metabolites and quantified as previously described, scaling back to the total ion chromatogram and normalizing to internal standard recovered, volume analyzed and mass extracted.

### Differential localization experiments

#### Plant inoculation and incubation

Combinations of barcoded strains were created by pelleting, washing, and resuspending glucose-grown overnight cultures (30 °C) in sterile, distilled water to OD 1, and then mixing them in equal amounts. *P. trichocarpa* BESC819, *P. tremula* x *P. alba* ‘INRA 717-1B4’, and the *PtrXB38-OE P. tremula* x *P. alba* ‘INRA 717-1B4’ were propagated according to previously published procedures [43]. In brief, sterile shoot tips were grown in tissue culture until root establishment (25 °C, 16 h photoperiod). Then, plants similar in size and development were chosen, 5 mL of microbe combination was mixed into 150 cm^3^ calcined clay for each plant, and the root systems were placed in the clay and gently buried. After an incubation time of 21-28 days in a closed system magenta box (same conditions as specified above), root systems were lifted from the clay, loosely attached clay particles removed, and the roots frozen at −20 °C until further processing. For differential localization analysis, 9 sampling locations were devised, consisting of a cross of three structural categories; primary, secondary, and tertiary, with three root regions; root tips, midzone, and root hairs (Figure 5A)[44].

#### Nucleic acid extraction

According to their development, roots were dissected into primary (oldest and thickest roots), secondary (first branching roots, medium size and age), and tertiary (youngest and thinnest). Furthermore, three sample types were collected for each structural category, i.e., tips (root tips of the structural category), hair roots (originating from the surface of the corresponding structure), and segments (root mass from the middle of the structure, whereby hair roots were removed from the surface). For each combination, 0.2 g of root mass was pooled for nucleic acid extraction and an initial pulverization step was conducted. This step consisted of freezing the tubes containing the root fragments in liquid nitrogen, and bead-beating them three times for 1 minute at 30 Hz using a TissueLyser II (Qiagen) and the steel beads included in the DNeasy Plant Pro Kit (Qiagen) with intermittent refreezing in liquid N_2_. Then, the pulverized root material was used as the regular input for said kit according to the manufacturer’s instructions. Bacterial mixtures used for inoculating plant experiments were extracted using the DNeasy Blood and Tissue Kit (Qiagen) according to the manufacturer’s recommendations.

### Library preparation, sequencing and data analysis

Microbial genomic DNA obtained from plant dissects and inoculum mixtures was amplified to enrich the barcode locus using primers suitable for sequencing adapter attachment depending on the intended sequencer (Table 4). Amplicons were then pooled at equimolar concentration and sequenced using an in-house Illumina MiSeq sequencer, as well as commercially via Illumina NovaSeq technology (VANTAGE, Vanderbilt University, Nashville, TN). From the resulting data, barcodes were extracted from sequencing reads by Bartender v1.1 [45], summarized, and then analyzed in R (v 4.0.3) using the tidyverse package (v 1.3.0) and a custom visualization script.

## Data availability

Raw sequencing reads generated and analyzed for this study can be downloaded from the NCBI Sequence Read Archive under Bioproject PRJNA1054559. Custom analysis scripts are available at https://github.com/s-christel/salicylate_barcode_experiment. All proteomics spectral data in this study was deposited at the ProteomeXchange Consortium via the MassIVErepository (https://massive.ucsd.edu/). The ProteomeXchange project identifier is PXD048223 and the MassIVE identifier is MSV000093756. The data can be reviewed under the username “MSV000093756_reviewer” and password “Christel_PMI”.

## Acknowledgements

This manuscript has been authored by UT-Battelle, LLC under Contract No. DE-AC05-00OR22725 with the U.S. Department of Energy. This work was supported by the Plant Microbe Interfaces Science Focus Area, funded by the Office of Biological and Environmental Research in the DOE Office of Science. The authors wish to acknowledge Dawn M. Klingeman for assistance with DNA amplicon sequencing.

## REFERENCES

1. Welch RA, Burland V, Plunkett G 3rd, Redford P, Roesch P, Rasko D, et al. Extensive mosaic structure revealed by the complete genome sequence of uropathogenic Escherichia coli. Proc Natl Acad Sci U S A 2002; 99: 17020– 17024.

2. Tettelin H, Masignani V, Cieslewicz MJ, Donati C, Medini D, Ward NL, et al. Genome analysis of multiple pathogenic isolates of *Streptococcus agalactiae*: Implications for the microbial ‘pan-genome’. Proceedings of the National Academy of Sciences 2005; 102: 13950–13955.

3. Niehus R, Mitri S, Fletcher AG, Foster KR. Migration and horizontal gene transfer divide microbial genomes into multiple niches. Nat Commun 2015; 6: 8924.

4. Goyal A. Metabolic adaptations underlying genome flexibility in prokaryotes. PLoS Genet 2018; 14: e1007763.

5. Pál C, Papp B, Lercher MJ. Adaptive evolution of bacterial metabolic networks by horizontal gene transfer. Nat Genet 2005; 37: 1372–1375.

6. Andreani NA, Hesse E, Vos M. Prokaryote genome fluidity is dependent on effective population size. ISME J 2017; 11: 1719–1721.

7. McInerney JO, McNally A, O’Connell MJ. Why prokaryotes have pangenomes. Nat Microbiol 2017; 2: 17040.

8. Baltrus DA. Exploring the costs of horizontal gene transfer. Trends Ecol Evol 2013; 28: 489–495.

9. Hall RJ, Whelan FJ, McInerney JO, Ou Y, Domingo-Sananes MR. Horizontal Gene Transfer as a Source of Conflict and Cooperation in Prokaryotes. Front Microbiol 2020; 11: 1569.

10. Karcagi I, Draskovits G, Umenhoffer K, Fekete G, Kovács K, Méhi O, et al. Indispensability of Horizontally Transferred Genes and Its Impact on Bacterial Genome Streamlining. Mol Biol Evol 2016; 33: 1257–1269.

11. Bruns H, Crüsemann M, Letzel A-C, Alanjary M, McInerney JO, Jensen PR, et al. Function-related replacement of bacterial siderophore pathways. ISME J 2018; 12: 320–329.

12. Hall JPJ, Wright RCT, Harrison E, Muddiman KJ, Wood AJ, Paterson S, et al. Plasmid fitness costs are caused by specific genetic conflicts enabling resolution by compensatory mutation. PLoS Biol 2021; 19: e3001225.

13. Michener JK, Vuilleumier S, Bringel F, Marx CJ. Phylogeny poorly predicts the utility of a challenging horizontally transferred gene in Methylobacterium strains. J Bacteriol 2014; 196: 2101–2107.

14. Shepherd ES, DeLoache WC, Pruss KM, Whitaker WR, Sonnenburg JL. An exclusive metabolic niche enables strain engraftment in the gut microbiota. Nature 2018; 557: 434–438.

15. Pudlo NA, Pereira GV, Parnami J, Cid M, Markert S, Tingley JP, et al. Diverse events have transferred genes for edible seaweed digestion from marine to human gut bacteria. Cell Host Microbe 2022; 30: 314–328.e11.

16. Major DW, McMaster ML, Cox EE, Edwards EA, Dworatzek SM, Hendrickson ER, et al. Field demonstration of successful bioaugmentation to achieve dechlorination of tetrachloroethene to ethene. Environ Sci Technol 2002; 36: 5106–5116.

17. Adrian L, Löffler FE. Outlook—The Next Frontiers for Research on Organohalide-Respiring Bacteria. In: Adrian L, Löffler FE (eds). Organohalide-Respiring Bacteria. 2016. Springer Berlin Heidelberg, Berlin, Heidelberg, pp 621–627.

18. Reinhold-Hurek B, Bünger W, Burbano CS, Sabale M, Hurek T. Roots shaping their microbiome: global hotspots for microbial activity. Annu Rev Phytopathol 2015; 53: 403–424.

19. Bais HP, Weir TL, Perry LG, Gilroy S, Vivanco JM. The role of root exudates in rhizosphere interactions with plants and other organisms. Annu Rev Plant Biol 2006; 57: 233–266.

20. Sun X, Xu Z, Xie J, Hesselberg-Thomsen V, Tan T, Zheng D, et al. Bacillus velezensis stimulates resident rhizosphere Pseudomonas stutzeri for plant health through metabolic interactions. ISME J 2022; 16: 774–787.

21. Dove NC, Veach AM, Muchero W, Wahl T, Stegen JC, Schadt CW, et al. Assembly of the Populus Microbiome Is Temporally Dynamic and Determined by Selective and Stochastic Factors. mSphere 2021; 6: e0131620.

22. Cregger MA, Carper DL, Christel S, Doktycz MJ, Labbé J, Michener JK, et al. Plant–Microbe Interactions: From Genes to Ecosystems Using Populus as a Model System. Phytobiomes Journal 2021; 5: 29–38.

23. Veach AM, Morris R, Yip DZ, Yang ZK, Engle NL, Cregger MA, et al. Rhizosphere microbiomes diverge among Populus trichocarpa plant-host genotypes and chemotypes, but it depends on soil origin. Microbiome 2019; 7: 76.

24. Boeckler GA, Gershenzon J, Unsicker SB. Phenolic glycosides of the Salicaceae and their role as anti-herbivore defenses. Phytochemistry 2011; 72: 1497–1509.

25. Dahal S, Hurst GB, Chourey K, Engle NL, Burdick LH, Morrell-Falvey JL, et al. Mechanism for Utilization of the Populus-Derived Metabolite Salicin by a Pseudomonas-Rahnella Co-Culture. Metabolites 2023; 13.

26. Lebeis SL, Paredes SH, Lundberg DS, Breakfield N, Gehring J, McDonald M, et al. PLANT MICROBIOME. Salicylic acid modulates colonization of the root microbiome by specific bacterial taxa. Science 2015; 349: 860–864.

27. Jun S-R, Wassenaar TM, Nookaew I, Hauser L, Wanchai V, Land M, et al. Diversity of Pseudomonas Genomes, Including Populus-Associated Isolates, as Revealed by Comparative Genome Analysis. Appl Environ Microbiol 2016; 82: 375–383.

28. Jiménez JI, Miñambres B, García JL, Díaz E. Genomic analysis of the aromatic catabolic pathways from Pseudomonas putida KT2440. Environ Microbiol 2002; 4: 824–841.

29. Carper DL, Weston DJ, Barde A, Timm CM, Lu T-Y, Burdick LH, et al. Cultivating the Bacterial Microbiota of Populus Roots. mSystems 2021; 6: e0130620.

30. Michener JK, Camargo Neves AA, Vuilleumier S, Bringel F, Marx CJ. Effective use of a horizontally-transferred pathway for dichloromethane catabolism requires post-transfer refinement. Elife 2014; 3.

31. Close DM, Cooper CJ, Wang X, Chirania P, Gupta M, Ossyra JR, et al. Horizontal transfer of a pathway for coumarate catabolism unexpectedly inhibits purine nucleotide biosynthesis. Mol Microbiol 2019; 112: 1784–1797.

32. Choi K-H, Schweizer HP. mini-Tn7 insertion in bacteria with single attTn7 sites: example Pseudomonas aeruginosa. Nat Protoc 2006; 1: 153–161.

33. Elmore JR, Furches A, Wolff GN, Gorday K, Guss AM. Development of a high efficiency integration system and promoter library for rapid modification of Pseudomonas putida KT2440. Metab Eng Commun 2017; 5: 1–8.

34. Elmore JR, Dexter GN, Baldino H, Huenemann JD, Francis R, Peabody GL 5th, et al. High-throughput genetic engineering of nonmodel and undomesticated bacteria via iterative site-specific genome integration. Sci Adv 2023; 9: eade1285.

35. Takahashi Y, Shintani M, Takase N, Kazo Y, Kawamura F, Hara H, et al. Modulation of primary cell function of host Pseudomonas bacteria by the conjugative plasmid pCAR1. Environ Microbiol 2015; 17: 134–155.

36. Nojiri H. Impact of catabolic plasmids on host cell physiology. Curr Opin Biotechnol 2013; 24: 423–430.

37. Yao T, Zhang J, Yates TB, Shrestha HK, Engle NL, Ployet R, et al. Expression quantitative trait loci mapping identified PtrXB38 as a key hub gene in adventitious root development in Populus. New Phytol 2023; 239: 2248–2264.

38. Choi K-H, Gaynor JB, White KG, Lopez C, Bosio CM, Karkhoff-Schweizer RR, et al. A Tn7-based broad-range bacterial cloning and expression system. Nat Methods 2005; 2: 443–448.

39. Chuanchuen R, Narasaki CT, Schweizer HP. Benchtop and microcentrifuge preparation of Pseudomonas aeruginosa competent cells. Biotechniques 2002; 33: 760, 762–3.

40. Batth TS, Tollenaere MX, Rüther P, Gonzalez-Franquesa A, Prabhakar BS, Bekker-Jensen S, et al. Protein Aggregation Capture on Microparticles Enables Multipurpose Proteomics Sample Preparation. Mol Cell Proteomics 2019; 18: 1027–1035.

41. Käll L, Canterbury JD, Weston J, Noble WS, MacCoss MJ. Semi-supervised learning for peptide identification from shotgun proteomics datasets. Nat Methods 2007; 4: 923–925.

42. Polpitiya AD, Qian W-J, Jaitly N, Petyuk VA, Adkins JN, Camp DG 2nd, et al. DAnTE: a statistical tool for quantitative analysis of −omics data. Bioinformatics 2008; 24: 1556–1558.

43. Henning JA, Weston DJ, Pelletier DA, Timm CM, Jawdy SS, Classen AT. Relatively rare root endophytic bacteria drive plant resource allocation patterns and tissue nutrient concentration in unpredictable ways. Am J Bot 2019; 106: 1423–1434.

44. Freschet GT, Pagès L, Iversen CM, Comas LH, Rewald B, Roumet C, et al. A starting guide to root ecology: strengthening ecological concepts and standardising root classification, sampling, processing and trait measurements. New Phytol 2021; 232: 973–1122.

45. Zhao L, Liu Z, Levy SF, Wu S. Bartender: a fast and accurate clustering algorithm to count barcode reads. Bioinformatics 2018; 34: 739–747.

