## Supplemental Figures for "Catabolic pathway acquisition by soil pseudomonads readily enables growth with salicyl alcohol but does not affect colonization of *Populus* roots"

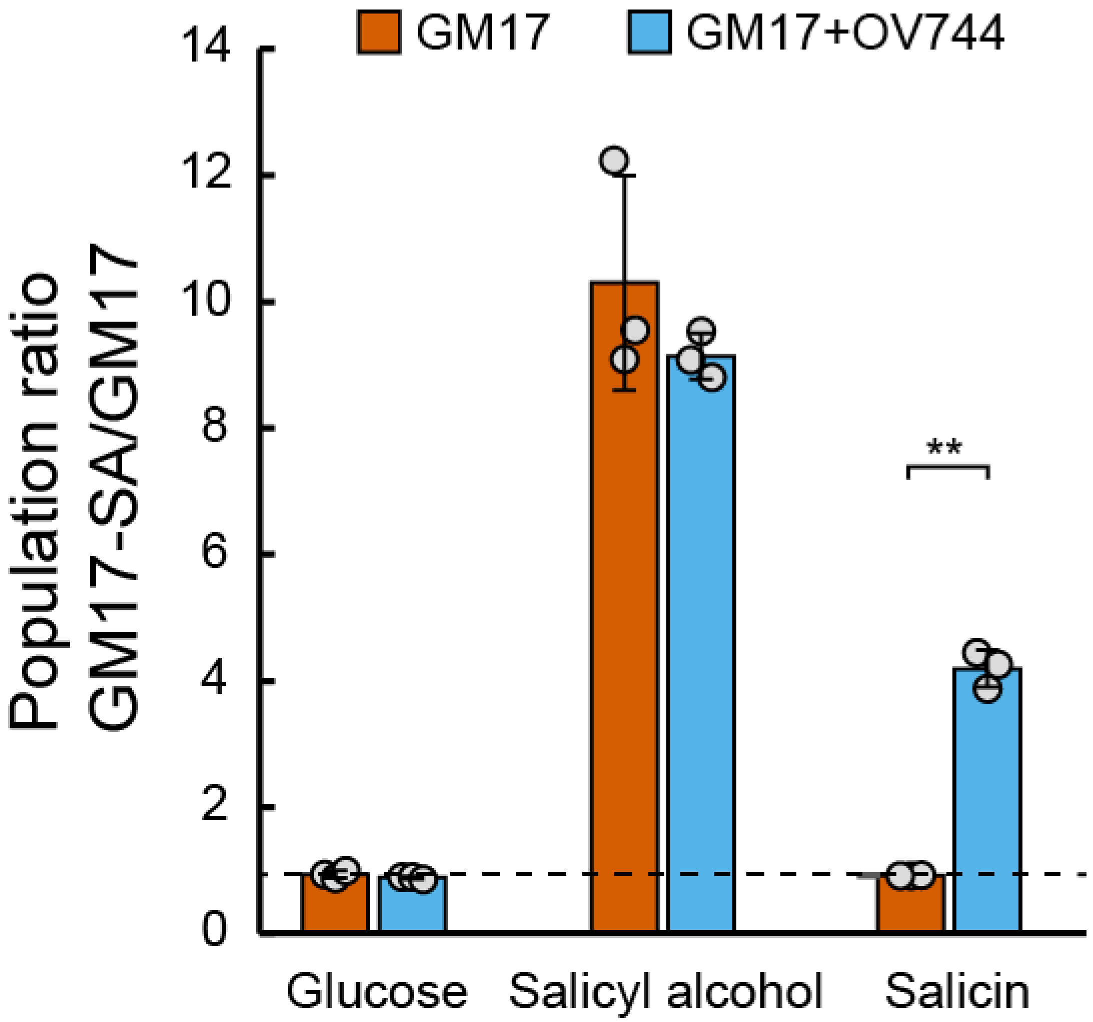


**Figure S1: The SA catabolic pathway provides an advantage during growth in liquid culture with SA**. A mixture of barcoded wild-type and engineered GM17 strains were grown in MOPS minimal medium with the indicated carbon source in the presence or absence of OV744. Population ratios before and after growth were calculated by barcode amplicon sequencing. The dashed line shows the population ratio for the inoculum. Error bars show one standard deviation, calculated from three biological replicates. **: p<0.01.

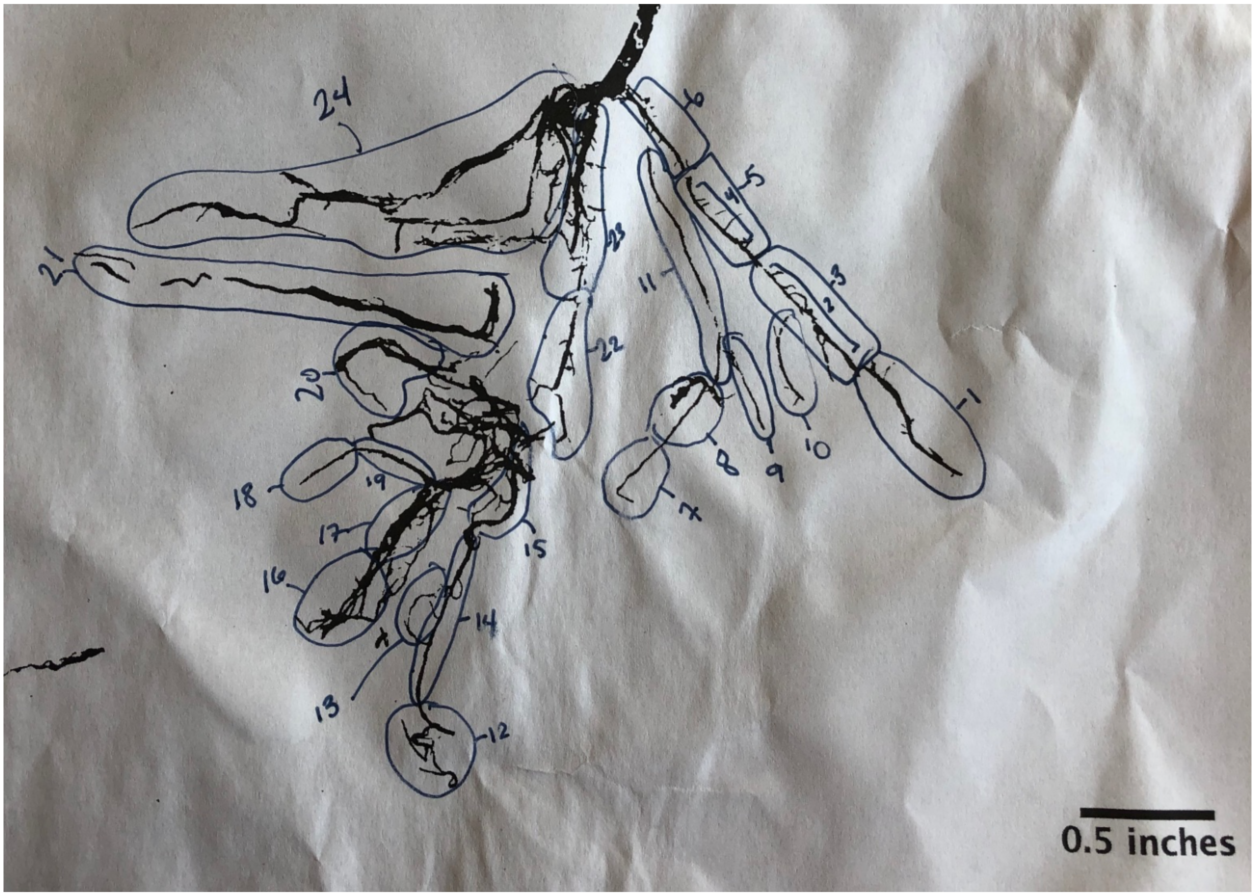


**Figure S2: Root dissection optimization tested a range of root sizes and orders.** A single tissue cultured *Populus trichocarpa* root system previously inoculated with barcoded wild-type *Pseudomonas* sp. GM17 was isolated, imaged, and dissected as indicated. Genomic DNA was extracted from each root segment and barcodes were amplified by PCR, as shown in Figure S3.

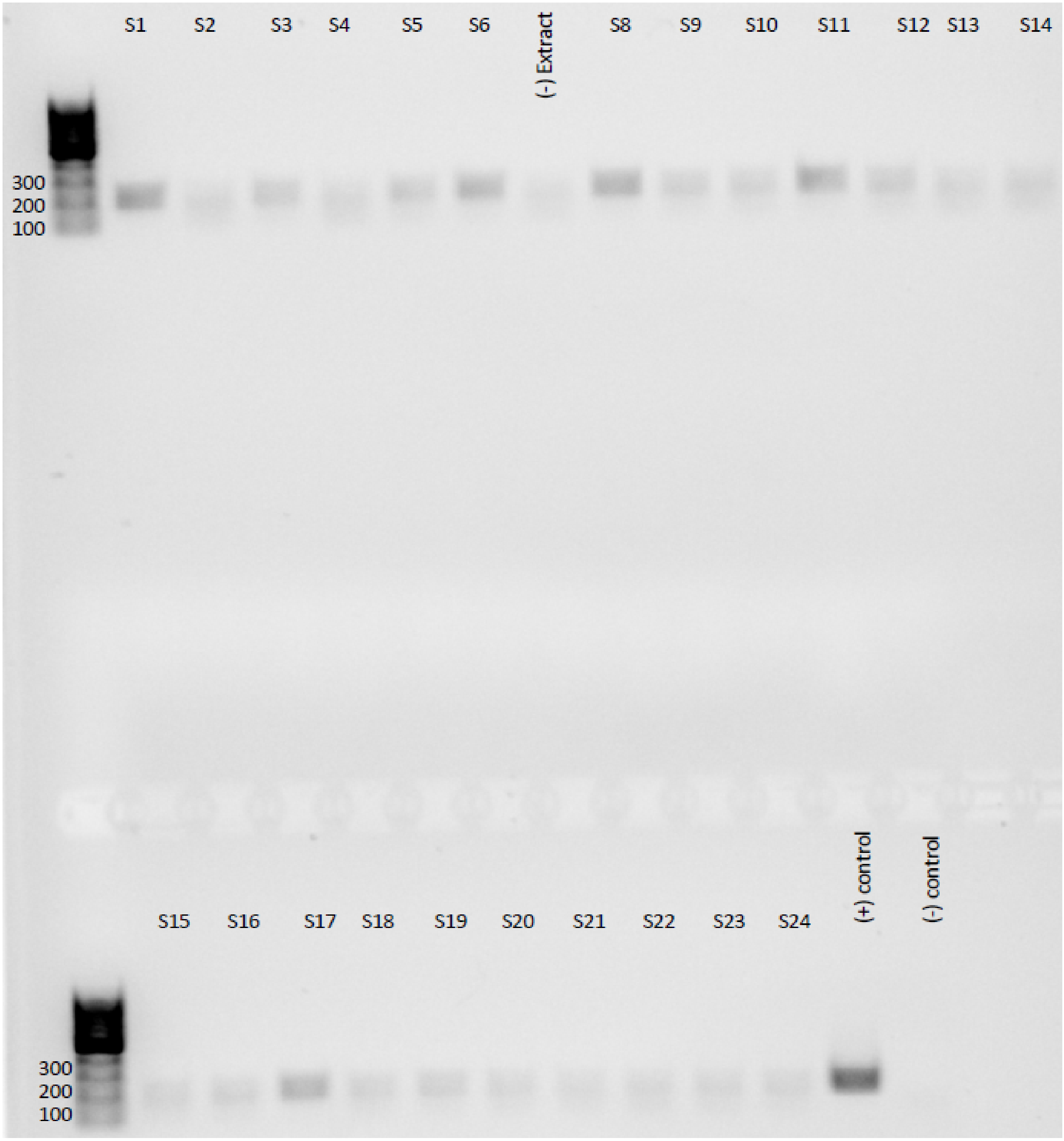


**Figure S3: Barcode amplification from root segments**. Sample IDs are as indicated in Figure S3. Barcodes were amplified by PCR and sequenced.

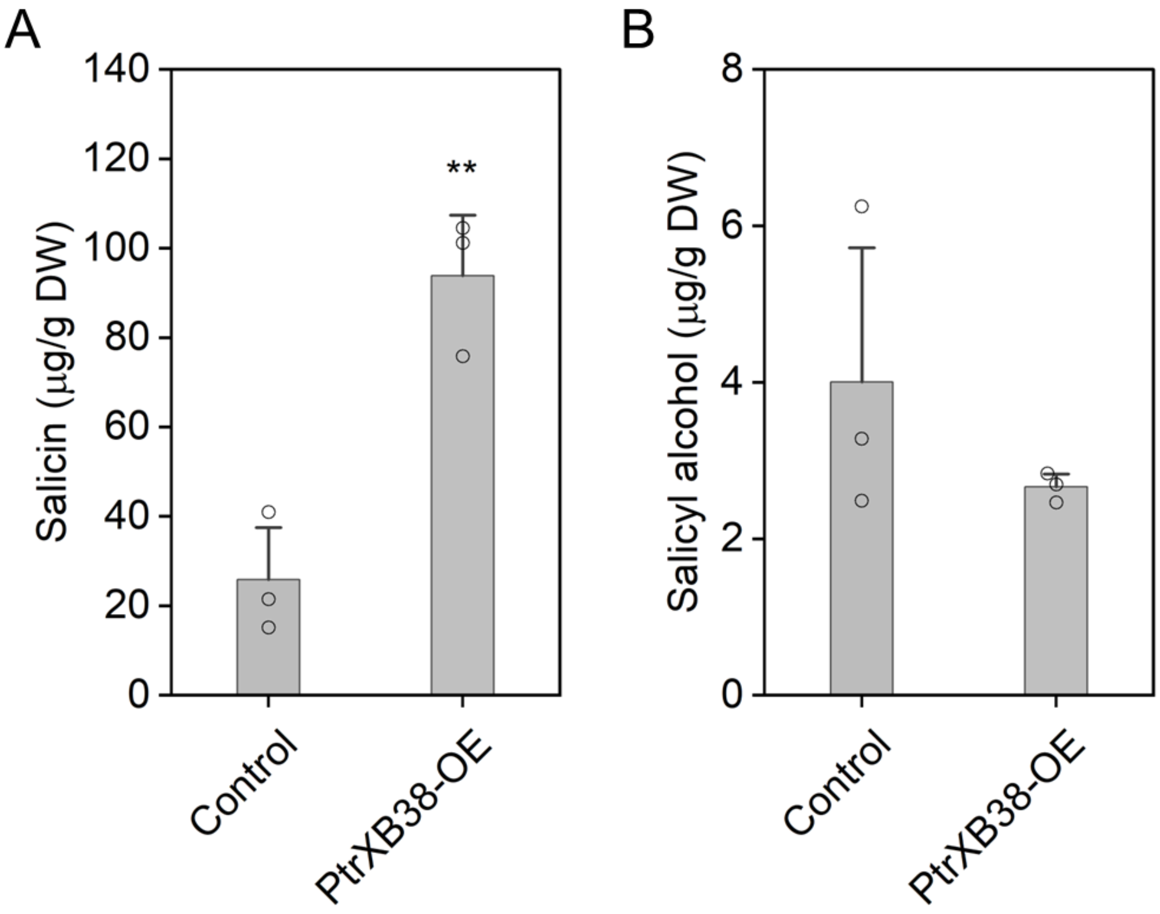


**Figure S4: Overexpression of PtrXB38 increases production of salicin in the roots of poplar (*Populus tremula* x *Populus alba*).** Roots from 2-month-old empty vector control and PtrXB38-OE transgenic plants were analyzed by GC-MS. Metabolite concentrations are relative to sorbitol, the internal standard. Bar charts represent mean ± SE (n= 3 independent plants), and double asterisk (**) represents significant difference between groups (P<0.01) by the Student's *t*-test.

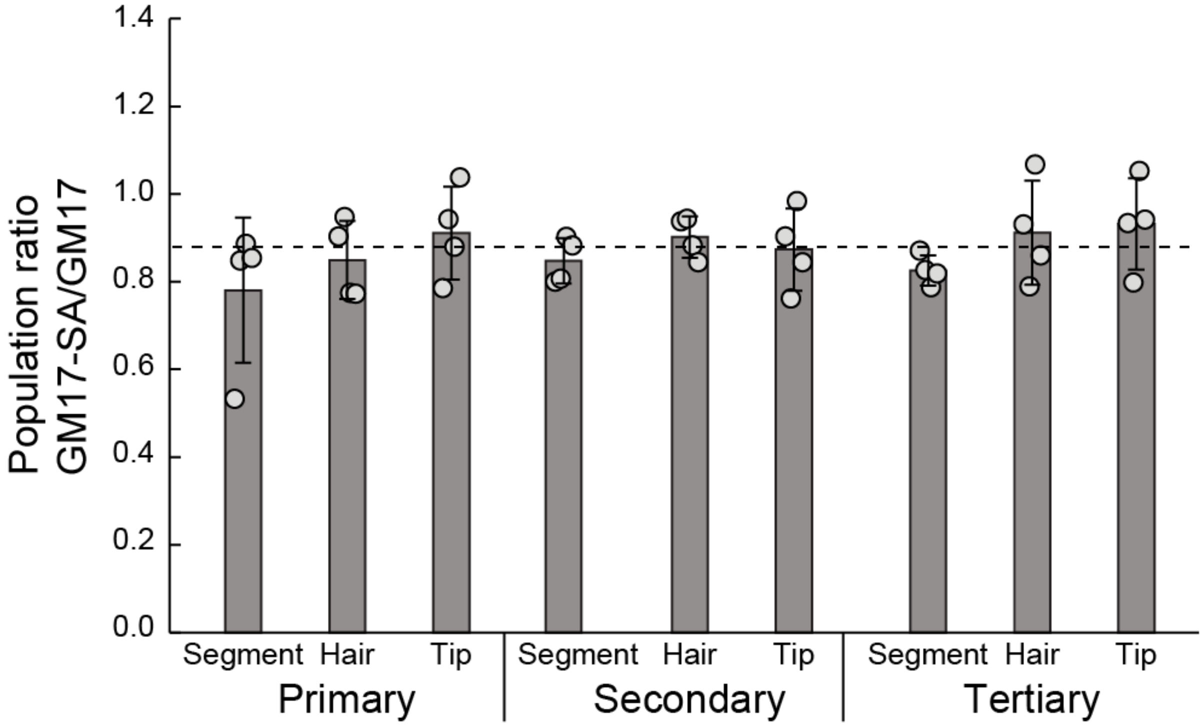


**Figure S5: SA catabolism does not provide a fitness advantage to GM17-SA during growth on XBAT35 trees.** Experiments were conducted as described in Figure 5D except without the addition of *Rahnella* sp. OV744.The dashed line shows the population ratio of the inoculum. Error bars show one standard deviation, calculated from the four biological replicates shown.
